# Optimizing error correction of RNAseq reads

**DOI:** 10.1101/020123

**Authors:** Matthew D. MacManes

## Abstract

**Motivation:** The correction of sequencing errors contained in Illumina reads derived from genomic DNA is a common pre-processing step in many *de novo* genome assembly pipelines, and has been shown to improved the quality of resultant assemblies. In contrast, the correction of errors in transcriptome sequence data is much less common, but can potentially yield similar improvements in mapping and assembly quality. This manuscript evaluates several popular read-correction tool’s ability to correct sequence errors commonplace to transcriptome derived Illumina reads.

**Results:** I evaluated the efficacy of correction of transcriptome derived sequencing reads using using several metrics across a variety of sequencing depths. This evaluation demonstrates a complex relationship between the quality of the correction, depth of sequencing, and hardware availability which results in variable recommendations depending on the goals of the experiment, tolerance for false positives, and depth of coverage. Overall, read error correction is an important step in read quality control, and should become a standard part of analytical pipelines.

**Availability:** Results are non-deterministically repeatable using AMI:ami-3dae4956 (MacManes_EC_2015) and the Makefile available here: https://goo.gl/oVIuE0

**Contact:** and @PeroMHC

## 1 INTRODUCTION

Genome-enabled biology – the study of biological phenomenon empowered by the use of high throughput sequencing of transcriptomes [MacManes and Eisen, 2014, Ferreira et al., 2013, Balakrishnan et al., 2014], genomes [Castoe et al., 2013, Bactrian Camels Genome Sequencing and Analysis Consortium et al., 2012], and epigenomes [Lyko et al., 2010, Lin et al., 2014] has grown in popularity over the past several years. Much of this growth has been driven by relatively cheap sequencing generated on the Illumina platform. Unlike the previous generation of sequence data (*e.g.,* Sanger) where error rates were far below 1%, the rate of error typical of the Illumina platform is between 1% and 3% [Wang et al., 2012]. This higher error rate is often considered mitigated by high sequencing coverage, which may often result in each nucleotide being sequenced more than 100 times (100x coverage). When depth of coverage the number or expected sequencing errors, these errors may be efficiently detected by assemblers and eliminated from the assembly, often early during the creation of the assembly graph [Compeau et al., 2011, Pevzner et al., 2001]. Though shotgun sequencing is expected produce uniform coverage, certain genomic features (*e.g.,* biased GC content) may inhibit the library construction or sequencing process, resulting in coverage valleys. The reconstruction of these regions may be significantly improved by read-error correction.

Concomitant with rapid improvement in quality and quantity of genomic data, and release of genome assemblers has been the development of novel algorithms for the correction of genomic data. These tools (reviewed in [Yang et al., 2013, Molnar and Ilie, 2014]) typically cluster identical and nearly-identical subreads of length k (*kmers*) then, in a probabilistic framework, attempt to minimize the Hamming (edit) distance of the reads to a consensus kmer. These algorithms can be brown down into four general classes (kmer-spectra, suffix array, multiple sequence alignment and Hidden Markov Model based methods) [Heo et al., 2014]. These algorithms assume uniform sequence coverage, and therefore have been applied most successfully to genomic data, where the have been shown to improve the quality of genome assembly [Salzberg et al., 2012].

In contrast to DNA sequencing of genomes, RNA sequencing of the expressed parts of the genome (*e.g.,* the transcriptome) offers unique challenges. Chief amongst these challenges include coverage that is variable with patterns of expression and the reconstruction of splice-isoforms, each of which may erode error correction algorithm’s ability to accurately distinguish sequencing error from meaningful variation. This complexity may result in higher than expected false-positive rate, or lower than expected rate of error correction. In the face of these challenges, error correction of transcriptome reads has been shown to improve transcriptome assembly [MacManes and Eisen, 2013]. Since it’s publication, new correction algorithms have been developed (*e.g.,* BLESS [Heo et al., 2014], and older methods have benefitted from new software implementations. In addition, newer evaluation algorithms had been developed (for example, see evaluation in Li, 2015). Given this, the current work aims to extend on previous work to additional error correction tools and newer analytics, while providing concrete recommendations to researchers interested in selecting an optimal software package for the correction of RNA sequencing reads.

## 2 METHODS

To evaluate the efficacy of read-based error correction of transcriptome data, I used a well characterized [Han et al., 2013] publicly available (SRR797058) Mus RNAseq dataset. Because the efficacy of error correction may vary with depth of sequencing, I randomly subsampled the full dataset to 10, 20, 50, 100 million paired end reads using the `subsampler.py` script available here (https://goo.gl/IfI3zm). The resultant subsets were trimmed with the software package Trimmomatic version 0.32 [Bolger et al., 2014] using recommendations from [MacManes, 2014]. Reads were then subjected to error correction using the following software packages: `SEECER` version 0.1.3 [Le et al., 2013], `Lighter` version 1.0.5 [Song et al., 2014], `SGA` version 0.10.13 [Simpson and Durbin, 2012], bfc version r177 [Li, 2015], and `BLESS` version 0.24 [Heo et al., 2014]. In correction algorithms (`SGA`, `BLESS`, `bfc`) that allowed for the use of larger *kmer* lengths, I elected to error correct with a small (*k* = 33) and a long (*k* = 55) *kmer*, while for the other software (`SEECER` and `Lighter`) that does not allow for longer *kmer* values, I set *k* = 31. `bfc` requires interleaved reads, which was accomplished using khmer version 1.3 [Brown et al., 2014].

After error correction, reads were mapped to chromosome 1 from the *Mus* genome (version GRCm38, available on Ensembl) using default settings of the software package bwa mem version 0.7.12-r1039 [Li, 2013]. The number of nucleotide mismatches between read and reference were calculated via the `nm` tag from the resultant SAM file. The difference in the number of mismatches between identically mapped reads between raw and error corrected reads was calculated using the `errstat.js` script contained in `bfc` and `K8`, contained in bwakit version 0.7.12 (https://github.com/lh3/bwa/tree/master/bwakit).

The required software has been installed on an Amazon EC2 machine image (AMI:ami-3dae4956). A makefile for recreating the analysis is located on GitHub (https://goo.gl/oVIuE0). Note the RAM requirements for determination of the appropriate size of instance.

## 3 RESULTS

Error correction of RNA sequencing resulted in a dramatic improvement in the number of error contains in sequence reads. This effect is highly variable depending on the specific error correction algorithm and *kmer* used as well as the depth of sequencing coverage. When reflecting on the different metrics, it becomes clear they vary with respect to the aggressiveness of correction. Researchers interested in selecting a tool may choose based on different metrics. For instance, some researchers may choose based on the corrector that makes the fewest mistakes while others may decide on a different optimality criteria. Despite the fact that the correctors are variable in efficacy, several patterns emerge. First, `SGA` is very aggressive when applied to transcriptome data. It makes, often by an order of magnitude, more reads better. This improvement however, if buffered by the fact that it makes the most reads worse and therefore is generally not an appropriate choice for the correction of transcriptome data. Next, the correction tool `Lighter` makes, in all of the tests conducted, the fewest number of reads worse (*e.g.,* increases the `nm` tag value infrequently). For researchers concerned about erroneous correction, this appears to be an optimal choice. Lastly, both correctors `bfc` and `Seecer` appear to preform well for a variety of metrics and all tested sequencing depths.

For low coverage transcriptome sequencing, the data presented in Tables 1 and 2 suggest that `bfc` may optimize error correction; this finding is constant in all of the tests involving lower coverage data. As the amount of sequencing coverage increases past 50 million paired end reads (Tables 3 and 4), the correction tool `Seecer` becomes more favorable, though this recommendation comes with the cautionary note on RAM usage. `Seecer` uses in excess of 1Gb of RAM per 1 million paired end reads. This reasonably large RAM requirement may be limiting for some researchers.

**Table 1.** 10 million paired end reads. `bfc` offers the best overall correction. RAM= the approximate amount of RAM required to complete error correction. lenkmer is the length of the *kmer*. Perfect is the number of reads that map perfectly (e.g., `nm:0`). Better is the number of reads whose `nm` flag is decreased after error correction relative to raw read mapping. Worse is the number of reads whose nm flag is increased after error correction relative to raw read mapping. Gain_perf is equal to (perfect - better) + better - worse. The error correction software */ kmer* length that maximizes (or minimizes in the case of worse) a given metric is indicated by bold-red type.

|  | RAM | kmer | perfect | better | worse | gain_perf |
| --- | --- | --- | --- | --- | --- | --- |
| raw |  |  | 7.17E+05 |  |  |  |
| bless |  |  |  |  |  |  |
|  | <16Gb | 33 | 9.88E+05 | 1.08E+06 | 8.20E+04 | 1.26E+06 |
|  | <16Gb | 55 | 9.40E+05 | 8.76E+05 | 4.59E+04 | 1.05E+06 |
| lighter |  |  |  |  |  |  |
|  | <16Gb | 31 | 9.74E+05 | 1.17E+06 | **4.27E+04** | 1.39E+06 |
| SGA |  |  |  |  |  |  |
|  | <16Gb | 33 | 7.65E+05 | **3.92E+06** | 3.81E+06 | 1.56E+05 |
|  | <16Gb | 55 | 7.73E+05 | 3.90E+06 | 3.82E+06 | 1.43E+05 |
| SEECER |  |  |  |  |  |  |
|  | <16Gb | 31 | 1.07E+06 | 1.24E+06 | 5.99E+04 | 1.54E+06 |
| bfc |  |  |  |  |  |  |
|  | <16Gb | 33 | **1.07E+06** | 1.48E+06 | 5.24E+04 | **1.78E+06** |
|  | <16Gb | 55 | 1.02E+06 | 1.22E+06 | 4.43E+04 | 1.48E+06 |

**Table 2.** 20 million paired end reads. `bfc` offers the best overall correction.

|  | RAM | kmer | perfect | better | worse | gain_perf |
| --- | --- | --- | --- | --- | --- | --- |
| raw |  |  | 1.43E+06 |  |  |  |
| bless |  |  |  |  |  |  |
|  | <16Gb | 33 | 2.01E+06 | 2.31E+06 | 1.98E+05 | 2.70E+06 |
|  | <16Gb | 55 | 1.90E+06 | 1.86E+06 | 9.99E+04 | 2.24E+06 |
| lighter |  |  |  |  |  |  |
|  | <16Gb | 31 | 1.88E+06 | 2.17E+06 | **8.31E+04** | 2.53E+06 |
| SGA |  |  |  |  |  |  |
|  | <16Gb | 33 | 1.49E+06 | **7.83E+06** | 7.63E+06 | 2.59E+05 |
|  | <16Gb | 55 | 1.54E+06 | 7.81E+06 | 7.63E+06 | 2.87E+05 |
| SEECER |  |  |  |  |  |  |
|  | ≈45Gb | 31 | **2.15E+06** | 2.52E+06 | 1.28E+05 | 3.12E+06 |
| bfc |  |  |  |  |  |  |
|  | <16Gb | 33 | 2.11E+06 | 2.86E+06 | 1.03E+05 | **3.44E+06** |
|  | <16Gb | 55 | 2.03E+06 | 2.44E+06 | 9.00E+04 | 2.96E+06 |

Same description as Table 1

**Table 3.** 50 million paired end reads. `SEECER` offers the best overall correction, with `bfc` close behind.

|  | RAM | kmer | perfect | better | worse | gain_perf |
| --- | --- | --- | --- | --- | --- | --- |
| raw |  |  | 3.58E+06 |  |  |  |
| bless |  |  |  |  |  |  |
|  | <16Gb | 33 | 5.15E+06 | 6.00E+06 | 6.30E+05 | 6.94E+06 |
|  | <16Gb | 55 | 4.89E+06 | 4.99E+06 | 3.12E+05 | 5.98E+06 |
| lighter |  |  |  |  |  |  |
|  | <16Gb | 31 | 4.64E+06 | 5.21E+06 | **2.03E+05** | 6.06E+06 |
| SGA |  |  |  |  |  |  |
|  | <16Gb | 33 | 3.62E+06 | **1.95E+07** | 1.91E+07 | 4.96E+05 |
|  | <16Gb | 55 | 3.74E+06 | 1.95E+07 | 1.91E+07 | 5.70E+05 |
| SEECER |  |  |  |  |  |  |
|  | ≈69Gb | 31 | **5.43E+06** | 6.44E+06 | 3.37E+05 | **7.95E+06** |
| bfc |  |  |  |  |  |  |
|  | <16Gb | 33 | 5.12E+06 | 6.63E+06 | 2.43E+05 | 7.93E+06 |
|  | <16Gb | 55 | 5.02E+06 | 5.96E+06 | 2.21E+05 | 7.18E+06 |

Same description as Table 1

**Table 4.**
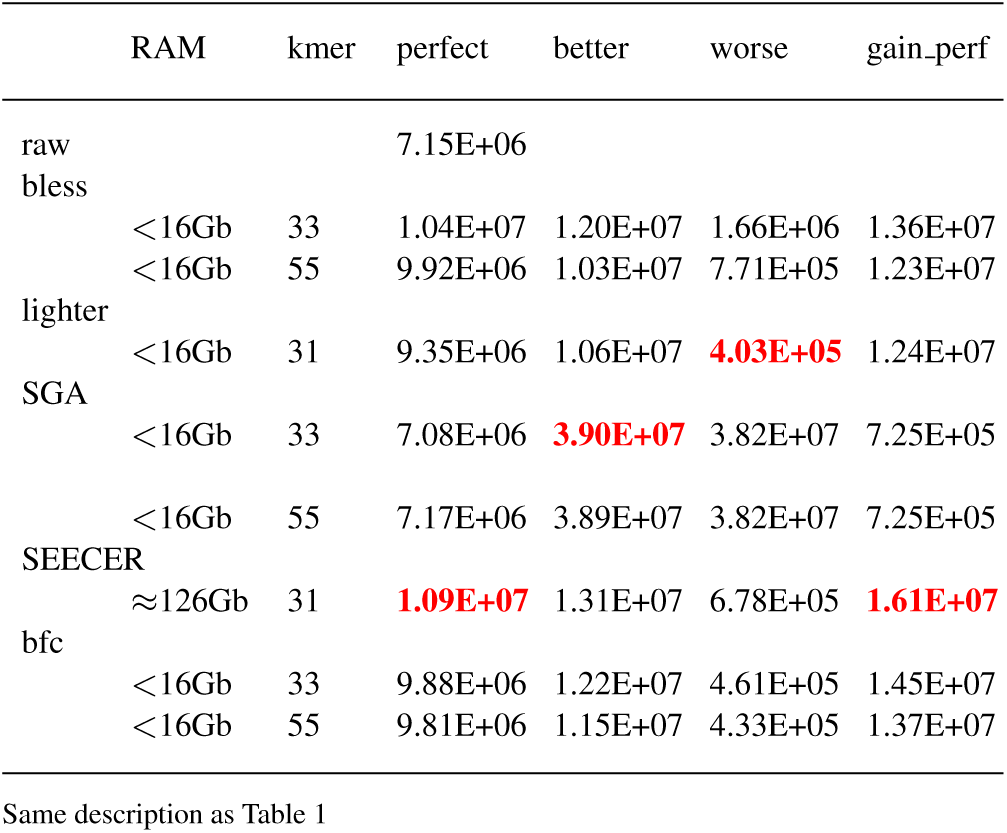
100 million paired end reads. `SEECER` offers the best overall correction.

## 4 DISCUSSION

`Lighter`, `bfc`, `SGA` and `BLESS` were all developed for genomic data – that many of them performed quite well was somewhat surprising, given the unique characteristics of transcriptome data. In contrast to these, `Seecer` was developed specifically for transcriptome sequence data. While it was most efficacious when applied to high coverage data (though with high RAM requirement), it also performed well for lower coverage datasets. bfc was most efficacious at low coverages, and was slightly worse than `Seecer` at higher coverage.

## 5 CONCLUSION

In conclusion, I offer the following recommendations for researchers interested in selecting an optimal tool for error correction.

1. For sequencing experiments where less than 50 million paired end reads are collected, the software `bfc` appears to offer an optimal solution, with `SEECER` running a close second.
2. For sequencing experiments where more than 50 million paired end reads are collected, the software `Seecer` is best, though at the cost of high RAM requirement. `Bfc` runs a close second.
3. In higher coverage data where a large amount of RAM is not available, `bfc` should be chosen.
4. If the research is sensitive to erroneous correction, even in the face of overall poorer performance, the `Lighter` package should be optimal.

## ACKNOWLEDGEMENT

*Funding*: This work was supported by start up funds provided by the College of Life Science and Agriculture at the University of New Hampshire.

